# Optimization of RNAi efficiency in PVD neuron of *C. elegans*

**DOI:** 10.1101/2023.08.31.555766

**Authors:** Pallavi Singh, Kavinila Selvarasu, Anindya Ghosh-Roy

**Author notes:** Correspondence: Anindya Ghosh-Roy, National Brain Research Centre, Manesar, NH-8, Nainwal Mode, Gurugram, Haryana-122052, India. These authors contributed equally (Pallavi Singh, Kavinila Selvarasu).

## Abstract

PVD neuron of *C. elegans* has become an attractive model for the study of dendrite development and regeneration due to the elaborate and stereotype dendrite morphology in this neuron. The molecular basis for dendrite maintenance and regeneration is poorly understood. RNA interference (RNAi) by feeding *E. coli* expressing dsRNA has been the basis of several genome wide screens performed using *C. elegans*. However, the feeding method often fails when it comes to nervous system. Using an optimal induction condition for the dsRNA expression in *E coli*, we fed the worm strains with HT115 bacteria expressing dsRNA against genes like *mec-3, hpo-30,* and *tiam-1*, whose loss of function are known to show dendrite morphology defects in PVD neuron. We found that RNAi of these genes in the strains such as *nre-1(-) lin-15b(-), lin-15b(-)* and *sid-1(-); lin-15b(-); Punc-119::sid-1[+]* resulted in significant reduction of dendrite branching. However, the phenotypes were significantly modest compared to the respective loss of function mutants. To obtain stronger phenotype for PVD specific genes, we have made a strain, which strongly expresses *sid-1* under *mec-3* promoter specific for PVD. When *Pmec-3::sid-1* is expressed in either *nre-1(-);lin-15b(-)* or *lin-15b(-)* background, the higher order branching phenotype after RNAi of *mec-3, hpo-30,* and *tiam-1* was significantly enhanced as compared to *nre-1(-);lin-15b(-)* and *lin-15b(-)* background alone. Next we tested the *nre-1(-) lin-15b(-),* P*mec-3-sid-1[+]* strain for the knockdown of genes playing role in dendrite regeneration process. We found that when *aff-1* and *ced-10* genes were knocked down in the *nre-1(-) lin-15b(-),* P*mec-3-sid-1[+]* background, the dendrite regeneration was significantly reduced and the extent of reduction was comparable to that of the mutants of *aff-1* and *ced-10*. Essentially, our strain expressing *sid-1* in PVD neuron optimizes the condition for RNAi for high throughput screening for PVD development, maintenance and regeneration.

## Introduction

PVD neuron in *C. elegans* is an excellent model to understand molecular basis of neuronal development and function (1). PVD neuron operates in multiple sensory modalities including harsh touch sensation and proprioception (2). Many concepts for neuronal polarization is understood using PVD neuron as a model (3, 4). PVD displays an orthogonal array of dendritic branches that cover major part of the body (1, 5). The higher order branches are arranged in a menorah like fashion (6). The tertiary branches from the adjacent menorahs self-avoid each other from physical contacts with the help of netrin *unc-40* signaling (7-9). *C. elegans* specific fusogen AFF-1 and EFF-1 sculpts its architecture (6). The guidance cue receptors and F-actin cytoskeleton machineries together help extending these branches (10, 11). The quaternary branches are stabilized by the physical contacts among epidermis and muscle (12, 13).

The anatomy of the PVD dendritic branches changes according to the mechanical stimuli (Inberg et al 2021). The dendrites show degeneration like phenomenon in older age due to abnormal expression of immunological peptides (14). Molecular control of these processes are poorly defined. It is also seen that upon laser mediated injury, the primary dendrites show various regeneration phenomena including the self-fusion between the proximal and distal dendrites, branching and regrowth (15-17). The self-fusion process is dependent on epidermal secretion of AFF-1 fusogen (16). The regrowth and branching is dependent on neuron specific function of CED-10 RAC GTPase (17). Dendrite regeneration phenomenon is recently also discovered both in fly and vertebrate model systems (18-20).

It might be relevant to investigate the conserved molecular mechanism controlling dendrite regeneration and degeneration in PVD neuron using high throughput screening. RNAi has been a powerful tool for knockdown of genes effectively to create loss of function condition (21, 22). This tool helps unbiased identification of the molecular pathways controlling a given biological process in worm system (23-25). However, it is often difficult to get effective knockdown of genes in the neurons using the feeding method (26). In order to overcome these problems, people have identified various sensitive strains which give enhanced RNAi (27-29). Particularly expressing the gene encoding RNA channel *sid-1* makes the neuron or any other tissue more sensitive to take up dsRNA (30-32). In this way, researchers have designed strains sensitive for RNAi in motor neurons and touch neuron (30, 33).

Although there are few studies which used RNAi to knockdown of candidate genes in PVD neuron, the penetrance of the phenotype seems to be low in these studies (7, 34, 35). There might be a scope to enhance the efficiency of RNAi in PVD with further modification of strains. In this study, we have first systematically analyzed the sensitized backgrounds for the loss of function phenotype for the genes such as *mec-3, hpo-30* and *tiam-1* that are required for dendritic branching in PVD. We found that the penetrance of the RNAi phenotype for a given gene in *nre-1(-) lin-15b(-)* or *sid-1(-) lin-15b(-); Punc-119::sid-1[+]* background is significantly lower as compared to the loss of function mutant allele of the corresponding gene. As a possible solution to improve the knockdown further, we have constructed a strain expressing the RNA channel SID-1 under the strong PVD-specific promoter *mec-3*. We found that when P*mec-3::sid-1* is expressed in *lin-15b(-)* or *nre-1(-) lin-15b(-)* background, it significantly enhances the phenotypes for the knockdown of *mec-3*, *hpo-30* and *tiam-1* as compared to the phenotypes seen in *lin-15b(-)* or *nre-1(-) lin-15b(-)* background alone. We further showed that the dendrite regeneration phenotype due to the RNAi mediated knockdown of *ced-10* and *aff-1* genes in *nre-1(-) lin-15b(-)*;P*mec-3*:: *sid-1[+]* background are comparable to the phenotype in the loss of function mutant alleles for the *ced-10* and *aff-1*. Therefore, combining the general RNAi sensitivity of the *nre-1(-) lin-15b(-)* background along with the PVD specific increase in *sid-1* level optimizes the RNAi efficiency in PVD neuron.

## Results

### RNAi mediated knockdown of PVD-specific genes related to dendritic branching does not cause penetrance of phenotype as close to loss of function mutants

Our goal was to find out the sensitive strain for the efficient knockdown of genes required for the function within the PVD neuron. In order to achieve this, first we wanted to optimize the efficient induction condition of ds-RNA production in the HT115 bacteria in our hand as different studies use a varying range of induction conditions for their experiments (22, 33, 36). In our hand, the induction of primary culture (condition-1) produced stronger phenotypes of genes such as *dhc-1*, *gbp-1*, and *unc-22* in wild type N2 Bristol strain as compared to the induction of secondary culture (condition-2) (Figure S1A). These genes and few others tested (Figure S1B) are ubiquitously required in worm and RNAi of these genes produce global phenotype in worm (36). Therefore, we used the condition-1 for pursuing the RNAi of the nervous system related genes whose knockdown produces severe uncoordinated or paralyzed phenotypes (Figure S1C). RNAi of *unc-14, unc-13, snb-1, unc-31* and *unc-25* revealed that both the *lin-15b(n744)* and *sid-1(-); lin-15b(-); Punc-119::sid-1(+)* strains produce consistent and high-penetrance loss of function phenotypes of these neuronal genes (Figure S1C). We next wanted to see whether any of these strains can knockdown genes known to shape the architecture of PVD dendrites (5), thereby producing relevant phenotypes. The dendrites in PVD neuron spans across the whole body (Figure 1A) and shows an orthogonal pattern in higher order branches (1°, 2°, 3° & 4^o,^ in Figure 1A). We selected *mec-3, hpo-30* and *tiam-1* which are required for the formation of these higher order branches in PVD (5, 37, 38). We performed RNAi on *lin-15b(-), nre-1(-) lin-15b(-)* and *sid-1(-);lin15b(-);Punc-119::sid-1(+)* strains and compared the penetrance of the branching phenotype with the null mutant of *mec-3* (Figure 1C-E). In *mec-3 (e1338)* mutant, the secondary / primary ratio per unit length of primary is closed to zero as there is no secondary branch in this mutant (Figure 1 B,C) as seen before (38). Whereas the RNAi of *mec-3* in the sensitive backgrounds caused a wide range of phenotypes (Figure 1B). Some population had partial quaternary (P4) or complete absence of quaternary (P4+3+2+1 or 3+2+1 in Figure 1B), in some both quaternary and tertiary missing (2+1 in Figure 1B). In rest of the population, the phenotype was like *mec-3* null mutant, where all the higher order branches were missing (Figure 1B). Therefore, the phenotype related to the secondary / primary ratio in the sensitive backgrounds due to the RNAi of *mec-3* gene was significantly weaker compared to the null mutant (Figure 1C). Likewise, both the quaternary/tertiary and tertiary/secondary ratios were significantly reduced in the sensitive backgrounds *lin-15b(-)* & *nre-1(-) lin-15b(-)* upon knockdown of *mec-3* gene (Figure 1D,E).

**Figure:1.**
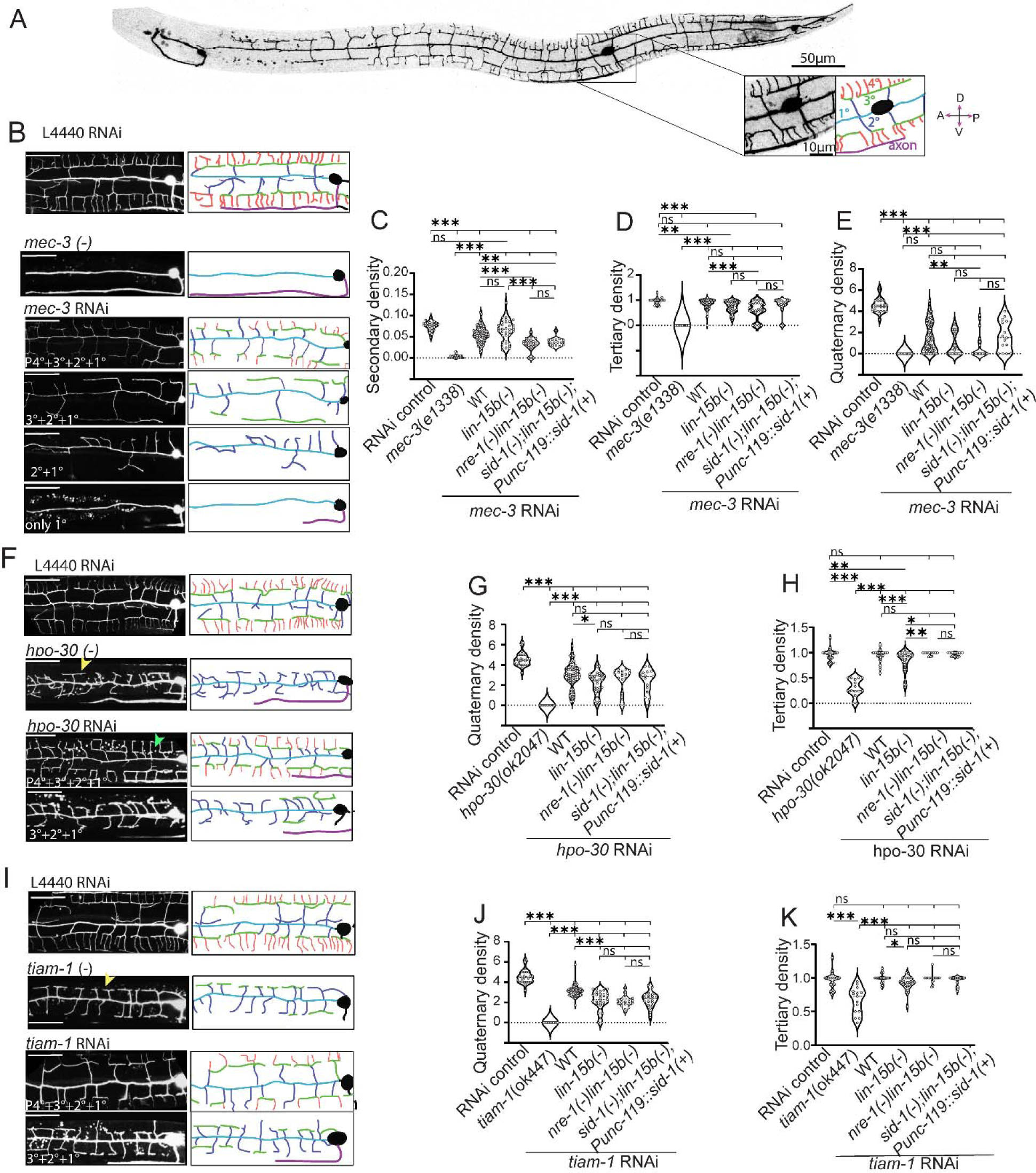
Standard sensitive strains cannot efficiently knockdown genes in PVD to cause dendritic phenotype close to the respective loss of function mutants. (A) The stitched confocal image of PVD neuron expressing GFP-reporter *wdIs52 [pF49H12.4:: GFP]* shows the elaborate dendritic branches of this neuron. The magnified inset and its schematic indicate the PVD soma, hierarchy of dendritic branches (1°/primary, 2°/secondary, 3°/Tertiary, and 4°/ quaternary) and its ventrally projecting axon. (B), (F), (I) Confocal images (left) and its schematic tracings (right) show the range of phenotype for dendritic branches shown in different colors, caused due to RNAi against PVD-specific genes such as *mec-3, hpo-30, tiam-1* in various sensitive background. The L4440 bacteria was used as RNAi control. The images of the respective loss of function mutants are also presented. The yellow arrowheads mark the tertiary with its missing quaternary branches in *hpo-30* and *tiam-1* mutant image. Green arrow shows the presence of partial quaternary in *hpo-30* RNAi worms. The plots (C-E) shows the density of secondary, tertiary and quaternary dendrite to give the quantitative description of *mec-3* RNAi and mutant phenotype. The plots (G-H) shows the density of quaternary, tertiary dendrite to demonstrate the *hpo-30* RNAi and mutant phenotype. Likewise, the plots (J-K) are the quaternary and tertiary density of *tiam-1* RNAi and mutant worms. Scale bar is 25 μm for all the images in (B),(F),(I). Statistics: For (C-E),(G-H),(J-K) One-way ANOVA with Turkey’s multiple comparison test, and number of worms (n), Biological replicates (N) are 15≤n≤67, 1≤N≤4, p<0.05*, 0.01**, 0.001***, ns, not significant.

The RNAi of many other genes that are known to control PVD morphology (5, {Smith, 2010 #9)} produced significant phenotype in *nre-1(-) lin-15b(-)* background using ‘condition-I’ for dsRNA induction in HT115 bacteria (Figure S2). Similarly, we tested the knockdown of genes like *hpo-30(ok2047)*, and *tiam-1(ok772)* required for the formation of the quaternary branches (10, 39). In the loss of function mutant of *hpo-30*, the quaternary/ tertiary ratio is closed to zero (Figure 1G) as the quaternary branches were completely missing in this mutant (yellow arrow/ Figure 1F).The knockdown of *hpo-30* using RNAi in the sensitive backgrounds also showed a reduction of quaternary branches (Figure 1F) although often some quaternary branches were still present (green arrow / Figure 1F). Therefore, the quaternary / tertiary ratio due to RNAi were significantly higher compared to the mutant (P<0.001,*** One-way ANOVA with Turkey’s multiple comparison test, Figure 1G). Similar trend was also noticed when we compared the quaternary phenotype in the loss of function mutant for *tiam-1* with the RNAi mediated knockdown of the same gene in the sensitive backgrounds (Figure 1-I,J).However, we were surprised to find that the strength of the phenotypes were comparable in *lin-15b(-)*, *nre-1(-) lin-15b(-)* and *sid-1(-);lin-15b(-);Punc-119::sid-1(+)* strains (Figure 1 C-E, G-H& J-K). We initially expected that *Punc-119::sid-1(+)* strain would cause significantly stronger penetrance as compared to the loss of either *lin-15b*(-) or *nre-1(-) lin-15b(-)* alone.

Therefore overall, our finding suggest that RNAi mediated knockdown of the genes in the available sensitive strains could not result in phenotype as close to loss of function mutants for the above-mentioned genes. This left the possibility for optimizing the RNAi efficiency in PVD neuron with further attention.

### PVD specific expression of *sid-1* helped knockdown of genes in PVD neuron

To enhance the phenotype for RNAi mediated knockdown of PVD-specific genes, we thought of utilizing the power of cell specific expression of RNA channel SID-1 (31, 32). The *sid-1* expression enhanced the phenotype of touch neuron and motor neuron by specific knockdown of genes using RNAi previously (30, 33). We looked for promoter that strongly expresses in PVD neuron using the neuronal cell atlas (CENGEN) (40, 41). We found that the expression of *unc-119* gene is relatively lower in PVD neuron (103.59 Transcripts per million/TPM). Whereas we noticed that *mec-3* is highly expressed in PVD (4067.391 TPM, CENGEN). Also, *mec-3* is expressed early in PVD, starting from early larval stages (7) allowing RNAi to begin early for targeted genes.

We made a single copy insertion of P*mec-3*::*sid-1*[+] (Figure 2A-C) by the mobilization of Mos1 transposon element (42, 43). As a control experiment we expressed P*mec-3*::GFP using the same single copy insertion transgenic method (42). The strain expressing P*mec-3*::GFP showed GFP expression in PVD (Arrowhead, Figure 2B) as well as in the PLM/ALM neuron (Figure 2B) validating the technique in our hand. The RNAi against GFP in Wildtype background causes loss of GFP expression in all the neurons representing the expression of the reporter promoter *wdIs52 [pF49H12.4:: GFP].* These are PVD, AQR and the tail neuron (Figure 2E). However, the RNAi against GFP in the strain expressing P*mec-3*::*sid-1* caused exclusive loss of GFP reporter in PVD neuron (Figure 2D-E) confirming that effect of *sid-1* is specific to PVD neuron. In 100% of the worms GFP was not visible in PVD neuron in P*mec-3*::*sid-1* background (Figure 2E). Similarly, when *hpo-30 and tiam-1* were knocked down using RNAi in the P*mec-3*::*sid-1* [+] strain, we found a reduction of quaternary branches (Figure 2 F-G), indicating the PVD specific effectiveness of the *sid-1* expression in the strain we developed.

**Figure 2:**
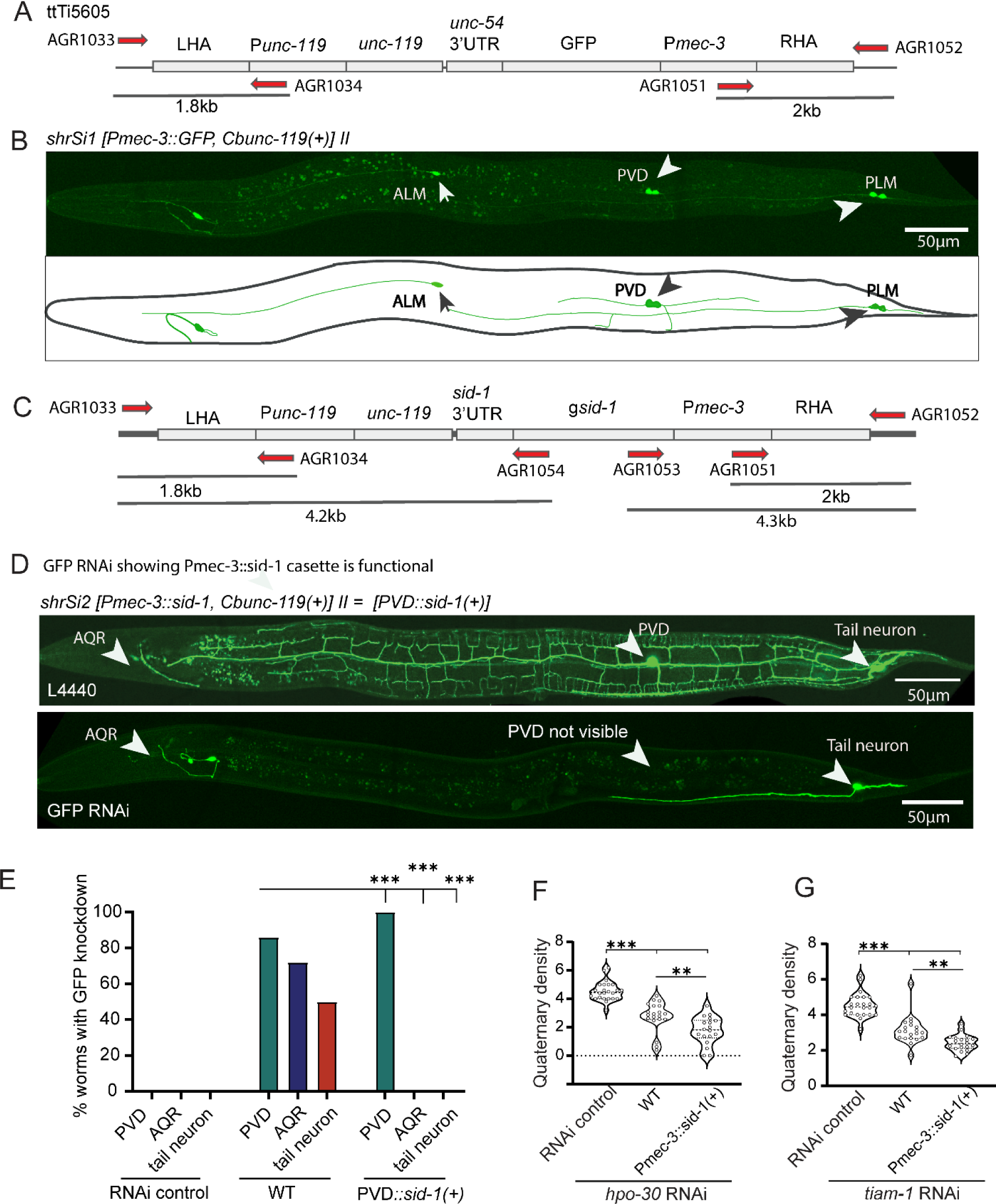
PVD specific expression of *sid-1* gives PVD-specific reduction of target-genes. (A) The illustration of P*mec-3::GFP* cassette inserted using MosI element ttTi5605 on chromosome II. LHA & RHA denotes Left Homology Arm and Right Homology Arm. (B) The confocal image and illustration showing the expression pattern of single copy inserted transgene *shrSI1*[*Pmec-3*::GFP]. The white arrowheads are marking GFP expression in PVD, ALM, and PLM neuron. (C) The illustration for the single copy insertion *shrSI2 [Pmec-3::sid-1]* driving *sid-1* in PVD neuron. (D) The confocal images of PVD neuron expressing *wdIs52 [pF49H12.4:: GFP]* after feeding the worms with L4440 (control) and GFP RNAi bacteria in *shrSI2 [Pmec-3::sid-1]* background. It shows specific reduction of GFP in PVD. White arrowheads shows three neurons AQR, PVD, Tail neuron visible in the *wdIs52 [pF49H12.4:: GFP]* reporter strain. (E) Percentage of worms with GFP knockdown in these three neurons are plotted separately for each genotypes showing PVD::*sid-1*(*Pmec-3*::*sid-1)* cassette effectively knocking down GFP in PVD neuron specifically. (F-G) Graph shows quaternary density for hpo-30 and tiam-1 RNAi fed worms was significantly reduced in *Pmec-3::sid-1(+)* strain. Statistics, For E, Fisher’s exact test 10≤n≤15, 1≤N≤2, (F-G) One-way ANOVA with Turkey’s multiple comparison test, 15≤n≤37, 1≤N≤2 p<0.05*, 0.01**, 0.001***, ns, not significant. Scale bar: 50μm. N stands for the number of independent replicates, and n stands for the number of worms taken.

### RNAi efficiency in PVD neuron can be synergistically enhanced by PVD specific expression of *sid-1* and sensitivity due to loss of *lin-15b*

Since the RNAi of *mec-3* gene in the existing sensitive strains such as *lin-15b(-)* or *nre1(-)lin-15b(-)* did not result in strong phenotypes like loss of function mutants, we hypothesized that elevating the level of *sid-1* in the sensitive strains might enhance the phenotypes in these strains. When we compared the secondary / primary ratio in the *nre-1(-)lin-15b(-)* and *nre1(-) lin-15b(-)*, P*mec-3::sid-1[+]* strains, we indeed found that adding *sid-1[+]* significantly enhanced the phenotype in the *nre-1(-) lin-15b(-)* (P< 0.01,** One-way ANOVA with Turkey’s multiple comparison test, Figure-3B). In many instances, in the *nre-1(-) lin-15b*, *Pmec-3::sid-1[+]* strain, only primary dendrite were visible (Figure 3A) as seen in *mec-3(e1338)* mutant. Adding *sid-1[+]* in the *lin-15b(-)* alone also significantly reduced the secondary / primary ratio (Figure 3A,B). Additionally, quaternary / tertiary ratio was also significantly dropped in the *nre-1(-) lin-15b(-)*, P*mec-3::sid-1[+]* as compared to *nre-1(-) lin-15b(-)* alone (Figure S3B).

**Figure 3:**
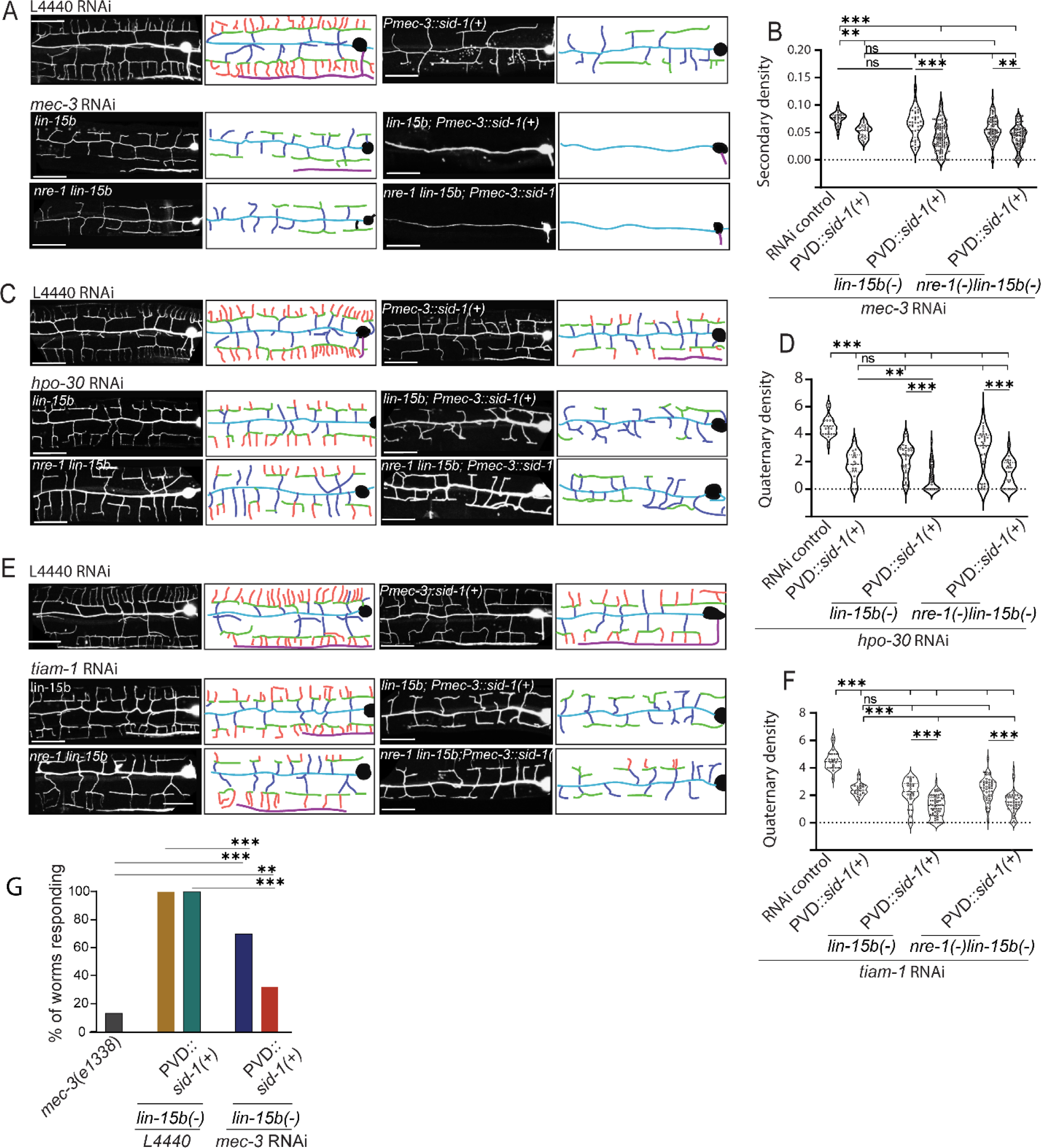
PVD specific expression of *sid-1* enhances the penetrance of RNAi phenotype. (A, C & E) The confocal images (left) and respective illustrations (right) show the RNAi phenotypes of *mec-3, hpo-30 or tiam-1* in standard sensitive backgrounds *lin-15b*(n744) or *nre-1(hd20) lin-15b(hd126)* and in strains with the addition of *Pmec-3::sid-1(+)* transgene in the similar sensitive backgrounds [*lin-15b*(n744) or *nre-1(hd20)lin-15b(hd126)]*. The *Pmec-3::sid-1(+)* transgene is denoted as *PVD::sid-1(+).* (B) The secondary density of PVD dendrite in the backgrounds with *mec-3* RNAi as shown in panel-A. An enhancement of phenotype was noticed in the strains having *PVD::sid-1(+)* transgene. Panel-D & F show the quaternary dendrite density in PVD upon the RNAi of *hpo-30* and *tiam-1*, respectively. (G) The percentage of worms responding to harsh touch delivered with a platinum wire in case of *mec-3* RNAi and *mec-3* mutant. The genotypes of the strains and the RNAi background are mentioned below the bars in the plot. showing mec-3 RNAi caused effective harsh touch defect in PVD specific RNAi sensitive background i.e *Pmec-3::sid-1*;*lin-15b*. Scale bar is 25μm. Statistics: For (B,D,F) One-way ANOVA with Turkey’s multiple comparison test, 24≤n≤97, 1≤N≤4, for (G) Fisher’s exact test, 10≤n≤20, 1≤N≤2 was performed, p<0.05*, 0.01**, 0.001***, ns, not significant, Biological replicates (N) and number of worms (n).

Similarly when we compared the quaternary / tertiary ratio in the *nre-1(-) lin-15b(-)* and *nre1(-) lin-15b(-)*, P*mec-3::sid-1[+]* strains in case of RNAi of *hpo-30*, we noticed that adding *sid-1[+]* significantly reduced the ratio in the *nre-1(-) lin-15b(-)* (P<0.001,*** One-way ANOVA with Turkey’s multiple comparison test, Figure 3C,D). Same trend was observed for the knockdown of *tiam-1* gene (P<0.001,*** One-way ANOVA with Turkey’s multiple comparison test, Figure 3E,F), which is exclusively required for the stabilization of the quaternary branches in PVD (10).

Since, adding *sid-1* transgene in the sensitive background produced loss of function phenotype for RNAi against *mec-3* for dendrite arbor, we also tested whether behavioral phenotypes caused by *mec-3* loss could also be obtained due to RNAi. We indeed found that upon RNAi of *mec-3*, there is a enhanced loss harsh touch response in *lin-15b(-)*,*Pmec-3::sid-1[+]* background as compared to *lin-15b(-)* single mutant (Figure 3G) as reported in *mec-3* mutant before (44).

Therefore, In both *lin-15b(-)*, *Pmec-3::sid-1[+]* and *nre1(-) lin-15b(-)*, *Pmec-3::sid-1[+]* strains we could achieve a synergistic effect of *lin-15b(-)* mutation and PVD specific expression of *sid-1* for the knockdown effect of PVD specific genes.

### The genes controlling dendrite regeneration pathways can be knocked down by RNAi in *nre1(-) lin-15b(-)*, *Pmec-3::sid-1[+]* background

Since the RNAi mediated knockdown of *mec-3, tiam-1* and *hpo-30* produced strong phenotype in *nre1(-) lin-15b(-)*, *Pmec-3::sid-1[+]* strain, we were encouraged to test whether one can optimally use this background in dendrite regeneration studies. Previous work showed that the primary dendrite of PVD upon laser injury shows regeneration response (15, 17). The dendrite regeneration depends on the RAC GTPase CED-10, GEF TIAM-1 (17) and the fusogen molecule AFF-1 (16). Typically, following dendrotomy (Orange laser shots, Figure 4A,), the primary dendrite regrows (green traces in schematic Figure 4B,C) and reconnects (green arrowheads) to the distal end of the injured dendrites (Figure 4B,C). The tertiary and quaternary branches corresponding to the menorahs of proximal and distal dendrites often are fused with each other (red highlighted boxes, Figure 4B) to bypass the gap created due to the injury. This phenomenon is called menorah-menorah fusion (16). As reported before (17), we found that extent of regrowth after dendrotomy as referred by ‘territory length’ is significantly reduced in the loss of function mutant of *ced-10* (Figure 4B). Similarly the percentage of reconnection and menorah-menorah fusion events were also significantly reduced in *ced-10* mutant (Figure 4E-F). Often there is a visible gap between the proximal and distal dendrites in the *ced-10* mutant (blue arrowhead, Figure 4B). When we performed RNAi of *ced-10* in *nre1(-)lin-15b(-)*, *Pmec-3::sid-1[+]* strain, we noticed all of these phenomena (blue arrowhead, Figure 4B) as seen in *ced-10* mutant. There were significant reduction in territory length, % reconnection and % menorah-menorah fusion on RNAi of *ced-10* (Figure 4D-F). More interestingly, the extent of reduction in the regeneration parameters were comparable in RNAi background and *ced-10* and *tiam-1*mutant (Figure 4D-F). Similarly, the reduction in regeneration parameters due to the RNAi of *aff-1* was comparable to the *aff-1* mutant (Figure 4B-F).

**Figure:4.**
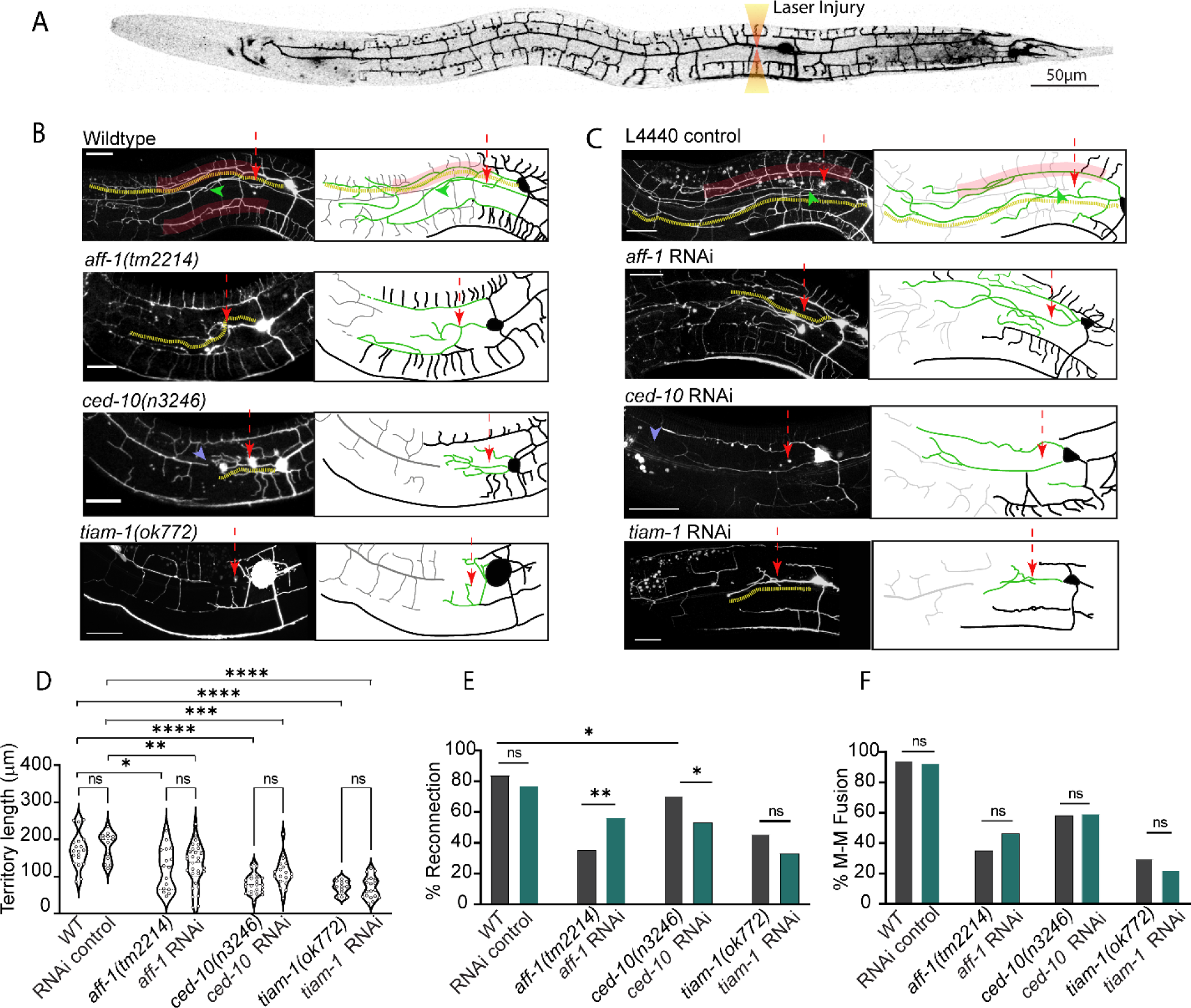
RNAi of genes required for PVD dendrite regeneration also showed phenotype enhancement in PVD specific *sid-1* expressing background. (A) Stitched confocal image of whole PVD showing the site chosen for Laser injury in dendrites i.e. close to first branch point (B) Images of regeneration pattern in mutant injured worms i.e. Wildtype (control), *aff-1(tm2214), ced-10(n3246), tiam-1(ok772)*. (C) Regenerating PVD dendrite in *Pmec-3::sid-1;nre-1(-)lin-15b(-)* RNAi worms fed *on L4440*(control)*, aff-1, ced-10, tiam-1* RNAi. In (B-C) confocal images(left) and schematics(right), menorah-menorah fusion is highlighted in red box, reconnection is marked with green arrowhead, yellow dotted line shows the extent of longest regenerating branch over images and schematic. Site of laser injury is marked with red arrows. Schematics on right side of these images have regrowing neurites shown in green traces, proximal PVD dendrite in black, distal portion after injury is drawn in grey, (D) Territory length i.e. length of longest neurite was measured and compared between mutants and RNAi of L4440(control), *aff-1, ced-10, tiam-1* genes in *nre-1(-)lin-15b(-)*;*Pmec-3::sid-1(+)* (E) Percentage of worms with reconnection events and (F) Percentage of worms with menorah-menorah fusion was compared for mutants and RNAi worms of these genes on *nre-1(-)lin-15b(-)*;*Pmec-3::sid-1(+)* strain. Statistics, For D, one-way ANOVA with Tukey’s multiple comparison test, 13≤n≤30, 1≤ N≤2, For (E-F) Fisher’s exact test were performed, 13≤n≤30, 1≤ N≤2, p<0.05*, 0.01**, 0.001***. ns, not significant. Scale bar for (B-C): 25μm. Biological replicates (N) and number of worms (n).

This suggested that the *nre-1(-) lin-15b(-)*, *Pmec-3::sid-1[+]* strain is quite efficient in the knockdown ability of various genes which are not only restricted to developmental branching but also required in adulthood for injury induced response.

## Discussion

PVD neuron has been a great model to understand the development and function of nerve cells (1, 2, 6, 45). Specially the stereotype and elaborate dendritic branches in PVD makes it an interesting system to explore the mechanism of dendrite development, maintenance and regeneration. Combining genetics and cell biology, people have made progress in mechanistic understanding of how elaborate anatomy of dendrite in PVD neuron is developed (6, 10). However, there is big gap in our understanding of how the dendritic arbor is maintained in adulthood and how it is repaired after injury.

RNAi has been instrumental in identifying novel molecular pathways in nerve cell development (29) and axon regeneration (46, 47) in worm model. However, RNAi is bit inefficient as well as variable depending on the neuron type and other conditions (48). Researchers always had to try various sensitive strains in order to get effective knockdown of genes in neurons (7, 29). The mutations in the genes such as *lin-15b*, *nre-1 lin15-b*, and *eri-1,* which negatively regulate RNAi process are often used to enhance the phenotypes in neuron (7, 29). The lack of expression of the RNA channel *sid-1* in the nervous system makes systemic RNAi inefficient in neuron (30). To overcome this challenge, researchers often mis-expressed *sid-1* in neuron to enhance the RNAi efficiency (30, 33). The existing strain that overexpresses *sid-1* under pan-neuronal promoter P*unc-119* did not give effective phenotype as compared to *nre-1(-) lin-15b(-)* or *lin-15b(-)* mutant alone. Therefore, we ventured our effort to try stronger PVD-specific promoter for obtaining stronger phenotype. Our results with the knockdown of *mec-3, hpo-30* and *tiam-1* genes indicated that indeed P*mec-3::sid-1* enhances the phenotype of dendrite branching in both the *lin-15b(-)* and *nre-1(-) lin-15b(-)* backgrounds. The *nre-1(-) lin-15b(-)* background also allows us to achieve the expected phenotype for dendrite regeneration following laser injury when *ced-10* and *aff-1* is knocked down. Therefore, this will be highly useful for the researchers studying the questions related to PVD neuron.

## Materials and Methods

### *C. elegans* strains and genetics

In this study, *C. elegans* strains were maintained at 20°C on OP50 bacterial lawn seeded over Nematode Growth Medium (NGM) plates (49). The loss of function mutation is represented as (-). For example, loss of function allele of *mec-3 (e1338)* represented as *mec-3(-).* The mutants used in this study are mostly loss of function by deletion or substitution unless otherwise mentioned. These mutants were obtained from Caenorhabditis Genetics Centre (CGC). The mutations crossed with *wdIs52 [pF49H12.4:: GFP]* strain carrying PVD specific GFP marker to aid visualization and microscopy and genotyped using their respective primers. Details of strain used for the study is provided in Table S1

### Optimization of the induction of ds RNA expression

To optimize the induction of dsRNA expression in HT115 bacteria in our hand, we tried three different induction conditions suggested in previous reports (22, 33, 36) with some modification i.e. condition 1 (primary culture), condition 2 (secondary culture with IPTG induction), condition 3 (secondary culture without IPTG induction). After seeding the bacteria grown under each condition, L4 staged worms (5-10 worms) were transferred onto these NGM plates containing carbenicillin, tetracycline, IPTG and was allowed to grow and give progenies to conduct experiments. The bacteria expressing various dsRNA were obtained from Arhinger’s and Vidal’s Library (50, 51), (Table S2)

### Condition I

The *E.coli* HT115 bacteria carrying RNAi clones targeting specific genes were thawed from -80deg and grown in Luria-Bertani (LB)-plates with 50 ug/ml carbenicillin and 12.5 ug/ml tetracycline and inoculated at 37^0^ C. Then a single colony was inoculated and grown at 37^0^ C in 4 ml LB in an incubator-shaker till it reached OD600 of 0.8. This primary culture was then pelleted and resuspended in 1X M9 buffer supplemented with 1.5 mM IPTG, carbenicillin and tetracycline. The resuspended culture was seeded onto NGM pates containing same concentration of carbenicillin, tetracycline, and IPTG. These plates were prepared two days in advance. Seeded plates were incubated at 25^0^ C for a duration of 36 hours for the induction of the RNAi construct within the bacteria. The condition-I involving the induction of primary culture was used before (36).

### Condition II

The primary culture was grown in LB medium containing 50 ug/ml carbenicillin and 12.5 ug/ml tetracycline at 37^0^ C overnight as described in ‘condition I’. The overnight grown primary culture then used to set secondary culture at 1 in 4 dilution in LB media containing carbenicillin, tetracycline, and 1Mm IPTG. The secondary culture was kept at 37^0^ C incubator-shaker until it reached an OD600 of 0.5-0.6. Subsequently the bacteria was pelleted down and resuspended in LB medium containing IPTG, carbenicillin, and tetracycline. Resuspended bacteria was seeded onto NGM plates containing antibiotics and IPTG. The plates were kept for induction period of 8 hours at 25^0^ C.

### Condition III

Same steps were performed as in Condition II with few modifications such as IPTG induction in secondary culture was not done and bacteria were grown until it reached OD600 of 0.5-0.6then, seeded plates were allowed to grow for 48hrs at room temperature as done before (33).

### RNAi using optimized condition

First optimal condition for induction of RNAi in *E. coli* was determined by performing RNAi against genes whose knockdown is known to produce strong phenotypes such as failure embryo-hatching, sterility or twitching of muscle etc. in N2 wild type strain. For example, we tested *dhc-1*, *gpb-1*, *unc-22* under three mentioned conditions in N2 background (Figure S1A). In our hand, the Condition-I produced stronger phenotypes as compared to the Condition-II and Condition-III (Figure S1A). Few other genes were also tested for further confirmation of the efficiency of ‘Condition-I’, such as *par-1, par-3, skn-1, dnc-1, bir-1, pal-1,, plk-1, ama-1,* which are ubiquitously required in worm and RNAi of these genes produce global phenotype (36). RNAi of many of the tested genes resulted in 100% penetrance in the wild type N2 Bristol strain (Figure S1B). Using this optimal induction condition (condition I), we performed RNAi of genes such as *unc-14, unc-13, snb-1, unc-31, unc-25* that are pan-neuronally required (30, 33), We used the strains that are shown to enhance RNAi sensitivity (Figure S1C).

### MOS1 element-related single copy insertion to construct P*mec-3::sid-1* transgenic strain

Mos1 transposon element-related method was used to insert P*mec-3::sid-1* in single copy as described before (42, 43). The P*mec-3::sid-1*+*unc-119* (pNBR65) was cloned in pcfj150 targeting vector (42) containing Mos-1 transposon element (Figure 2A). The genomic *sid-1* was amplified from *sid-1* plasmid (TU866) (30) using AGR1016 and AGR1017 primers. The pcfj150 backbone was amplified using AGR 1012 and AGR1013 primers and *mec-3* promoter was amplified with primers AGR1014 and AGR1015. These three fragments were assembled into a plasmid using In-Fusion reaction (cloning primer details in Table S3). For the confirmation of expression pattern of *mec-3* promoter, *Pmec-3::GFP+unc-119* (pNBR64 cassette was made (Figure 2A).

The uncordinated progenies of EG6699 [ttTi5605 II; *unc-119(ed9)*III; oxEx1578 [eft-3p::GFP+ Cbr-unc-119(+)] strain were injected with injection mix containing plasmids 50ng/ul Pcfj601(P*eft-3*::transposase), 50ng/ul of cassette *Pmec-3::sid-1+unc-119*, 29ng/ul Pma122 (Phsp::*peel-1*) along with marker plasmids i.e. 10ng/ul Pgh8 (*Prab-3*::mcherry), 2.5ng/ul pcfj90 (P*myo-2*::mcherry), 5ng/ul pcfj104 (*Pmyo-3*::mcherry), 20ng/ul 100 bp ladder. First, for confirmation of expression pattern of *mec-3* promoter, *Pmec-3*::GFP+*unc-119* cassette was also inserted using same strategy (Frøkjær-Jensen et al 2012). Around 60-70 worms were injected and two worms were kept in each plate. Plates were kept at 25^0^ C for a week till starved. Then for 3 hours it was kept at 34^0^ C for *Peel-1* mediated negative selection, worms with extrachromosomal array can’t survive this heat shock. Then, these plates were transferred to 20^0^ C for a day, 10 healthy L2-L3 non-unc worms were chosen from plates and genotyped using PCR, made homozygous for *Pmec-3::sid-1*(+) and outcrossed. For confirmation of insertion two PCR of 1.8kb and 2kb was done using primers (AGR1033 & AGR1034, Table S3) on upstream of ttTi5605 and Punc-119 respectively (Figure 2). Other set of primers (AGR1051 & AGR1052, Table S3) were designed on other end i.e downstream to ttTi5605 and *Pmec-3* (Figure 2). For confirmation of full *Pmec-3::sid-1* cassette insertion, long PCR of 4.2kb and 4.3kb using primers AGR1053 and AGR1054 was also performed (Figure 2). PCR primer details are given in Table S3. After confirmation by PCR and outcrossing, we used *shrSI1* [*Pmec-3::*GFP*+unc-119*] and *shrSI2* [*Pmec-3:: sid-1* + *unc-119*], for our experiments as shown in Figure-2, 3 and 4.

### Imaging of PVD neuron

The worms were mounted in 10 mM Levamisole hydrochloride (Sigma®) solution on the 5% agarose (Sigma®) pads made on the glass-slides. The worms were imaged with 63X/1.4NA oil objective of Nikon® A1R confocal system at a voxel resolution of 0.41µm x 0.41µm x 1µm and tile imaging module using imaging lasers 488nm(GFP), 543nm (mCherry/RFP) with 1-1.8 AU pinhole at 512x512 pixel resolution files for further analysis. For regeneration study, images were obtained at 24-hrs post-injury using same imaging condition.

### Dendrite branch quantification

PVD dendrite branch density was quantified as quaternary density, tertiary density and secondary density to normalize the phenotypes acquired in images (Figure 1b,F & I) encompassing from cell body till the middle of major dendrite using following formula: Quaternary density: Total number of quaternary dendrites / Total number of tertiary dendrites. Tertiary density: Total number of tertiary dendrites/ Total number of secondary dendrites. Secondary density: Total number of secondary dendrites / length of primary dendrite(µm). The length of primary dendrite was measured using Simple Neurite Tracer plugin in Fiji-ImageJ®.

### Laser system and dendrotomy details

Dendrotomy were conducted on worms at the L4stage using the Bruker® ULTIMA system with spectraPhysics® Two-photon femtosecond laser. This laser is tunable and operates in the infrared range (690-1040 nm). The laser output was controlled using Conoptics pockel cells. For visualization of the PVD and injury, lasers with wavelengths of 920nm and 720nm were used simultaneously by two sets of galvanometer mirror scanning X-Y (Brar et al 2022). To prepare slides, worms were immobilized using Levamisole hydrochloride (10mM) on 5% agarose pads and mounted with Corning cover glass. Worm-containing slides were placed under 60X/0.9NA water objective (Olympus®) with a pixel resolution of 0.29um x 0.29um.

During the experiments (Figure. 4), the PVD dendrites were severed at the first branch point, approximately 10 um away from the cell body, using the first laser shot. This was followed by one more consecutive shot with a relative distance of 10-15um from the previous shot, resulting visible gap. After injury, worms were transferred to freshly seeded NGM plates with OP50 or RNAi bacteria for further observation.

### Dendrite Regeneration Quantification

Dendrite regeneration was quantified based on regrowth from site of injury and fusion related parameters like menorah-menorah fusion (15, 16) where one big menorah is supported by more than one secondary branch (Figure.4B-C,red highlighted box). Also we see these regrowing neurites getting connected to distal dendrite (Figure.4B-C, green arrowhead) evaluated as reconnection events (17). The extent of territory covered by regrowing dendrite (Figure 4B-C, yellow dotted line) was measured using Simple Neurite Tracer plugin in Fiji-ImageJ® tracking the longest regenerating dendrite from cell body to the end point.

### Harsh Touch behavior analysis

The worms fed on RNAi bacteria such as L4440 (empty vector/Control) and *mec-3* (dsRNA against *mec-3*) were single-selfed in 10-20 plates using eye lash pick and left for few minutes. Videos were recorded while giving them harsh touch with platinum wire posterior to vulva, recording was done for nearly 20-30 seconds as described before (2, 52). Percentage of worms with positive response were calculated which showed observable increase in speed indicating escape response after harsh touch as represented in Figure 3G.

## Statistical analysis

The statistical analysis were performed using GraphPad Prism software (version 9.5.1). For two-sample comparisons, unpaired two tailed t-tests were used. When analyzing multiple samples, one way analysis of variance (ANOVA) was performed, followed by Tukey’s multiple comparisons test. The data used for the ANOVA analysis consisted of naturally occurring data with a normal distribution spread. To compare population data, fraction values were calculated for each sample and compared using a two-tailed chi-square Fischer’s exact contingency test.

The figure legends of the respective bar provides the information about the number of samples (n) and the number of biological replicates (N). The significance levels considered for all statistical experiments were p<0.05*, 0.01**, 0.001***.

wdIs52wdIs52

## Acknowledgments

We thank Yuji Kohara for cDNAs. We thank National BioResource Project (NBRP), Japan, and Caenorhabditis Genetics Center (CGC) for strains. CGC is supported by the NIH Office of Research Infrastructure Programs (P40 OD010440). This work is supported by the NBRC core fund from the Department of Biotechnology, DBT/Wellcome Trust India Alliance (Grant # IA/I/13/1/500874 to A.G.-R.) and a grant from the Science and Engineering Research Board (SERB: CRG/2019/002194 to A.G.-R.). We thank Martin Chalfie for gifting the *sid-1* transgene (TU866), Erik Jorgenson for gifting the plasmid reagents for generating Mos1 insertion strain and Santosh Kumar for the advice in making the injection mix. We also thank Arnab Mukhopadhyay and Sachin Kotak for various RNAi clones.

## Author Contributions

P. Singh, K. Selvarasu, and A. Ghosh-Roy designed experiments. P. Singh, and Kavinila S, performed experiments and analyzed data. K. Selvarasu, P. Singh and A. Ghosh-Roy wrote the manuscript.

