## Supporting File for "Optimization of RNAi efficiency in PVD neuron of *C. elegans*"

**List of supplementary figures**

**Figure S1, related to Figure 1**

**Figure S2, related to Figure 1**

**Figure S3, related to Figure 3**

**List of tables**

**Table S1**  List of *C. elegans* strains used in this paper.

**Table S2** List of RNAi bacteria used in this paper.

**Table S3** List of primers used for the cloning, sequencing and genotyping purposes.

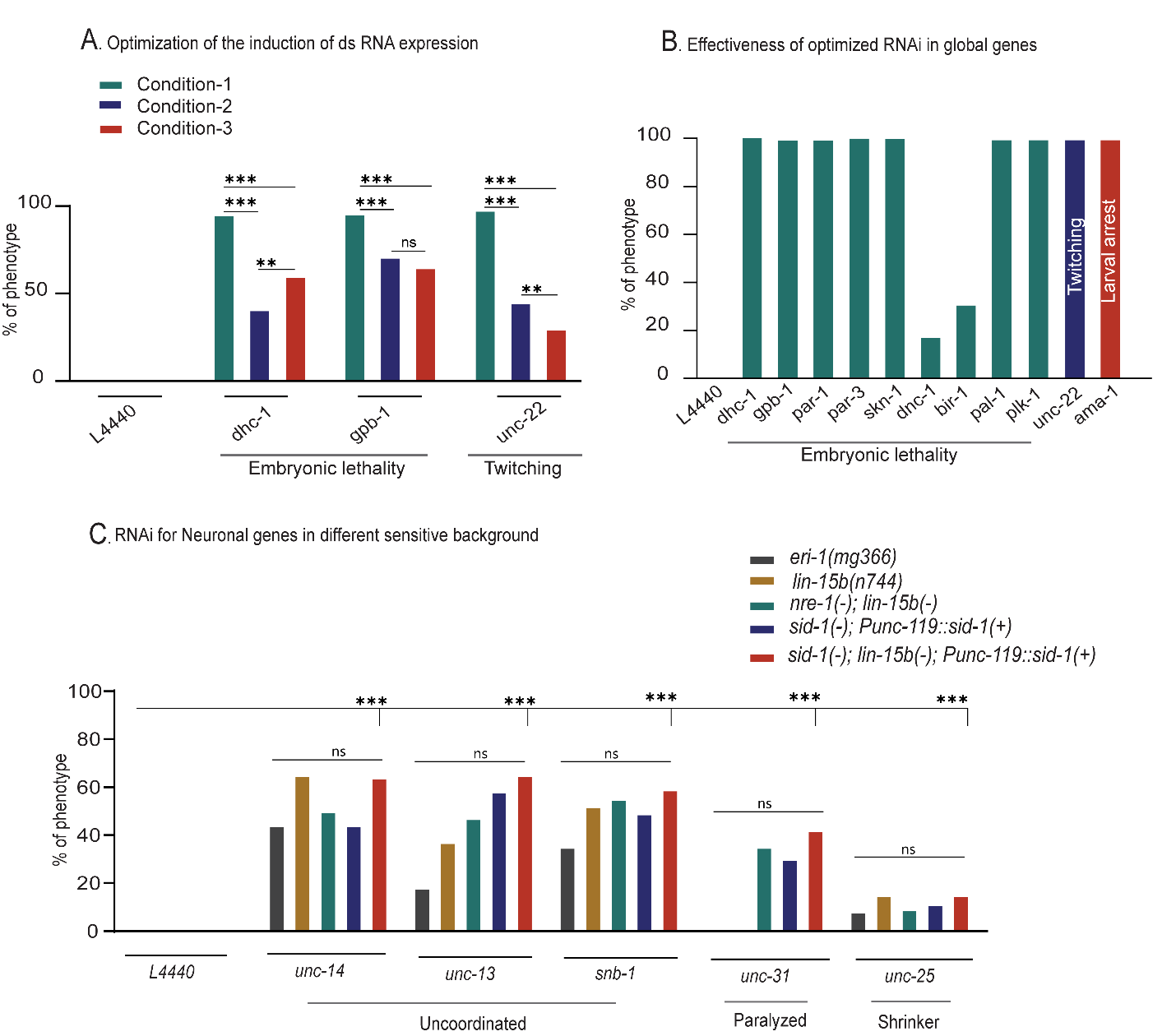

**Figure S1, related to Figure 1:**

(A) The percentage of phenotype involving embryonic lethality and twitching in L4440 (control), *dhc-1, gbp-1, unc-22* RNAi done in N2 Bristol background under three different induction conditions are plotted. Condition-I: induction of primary culture, Condition-II: induction of secondary culture with IPTG, and Condition-III: secondary culture was grown without IPTG induction. (B) The effectiveness of RNAi in N2 background using “condition-I” was further verified by knocking down various genes that cause embryonic lethality, twitching, larval arrest. (C) The organism-level phenotypes caused due to RNAi for genes required pan-neuronally are shown in this bar-plot. In this experiment, the RNAi was performed in the sensitive genetic backgrounds such as *eri-1(mg366),* *lin-15b(n744),* *nre-1(hd20)lin-15b(hd126)* and neuronal sensitive background *sid-1(pk3321);Punc-119::sid-1(+)* and *sid-1(pk3321);lin-15b(n744);Punc-119::sid-1(+)*. Biological replicates (1≤N≤2). (A-C) Statistics: Fisher’s exact test were performed, p<0.05*, 0.01**, 0.001***, ns, not significant.

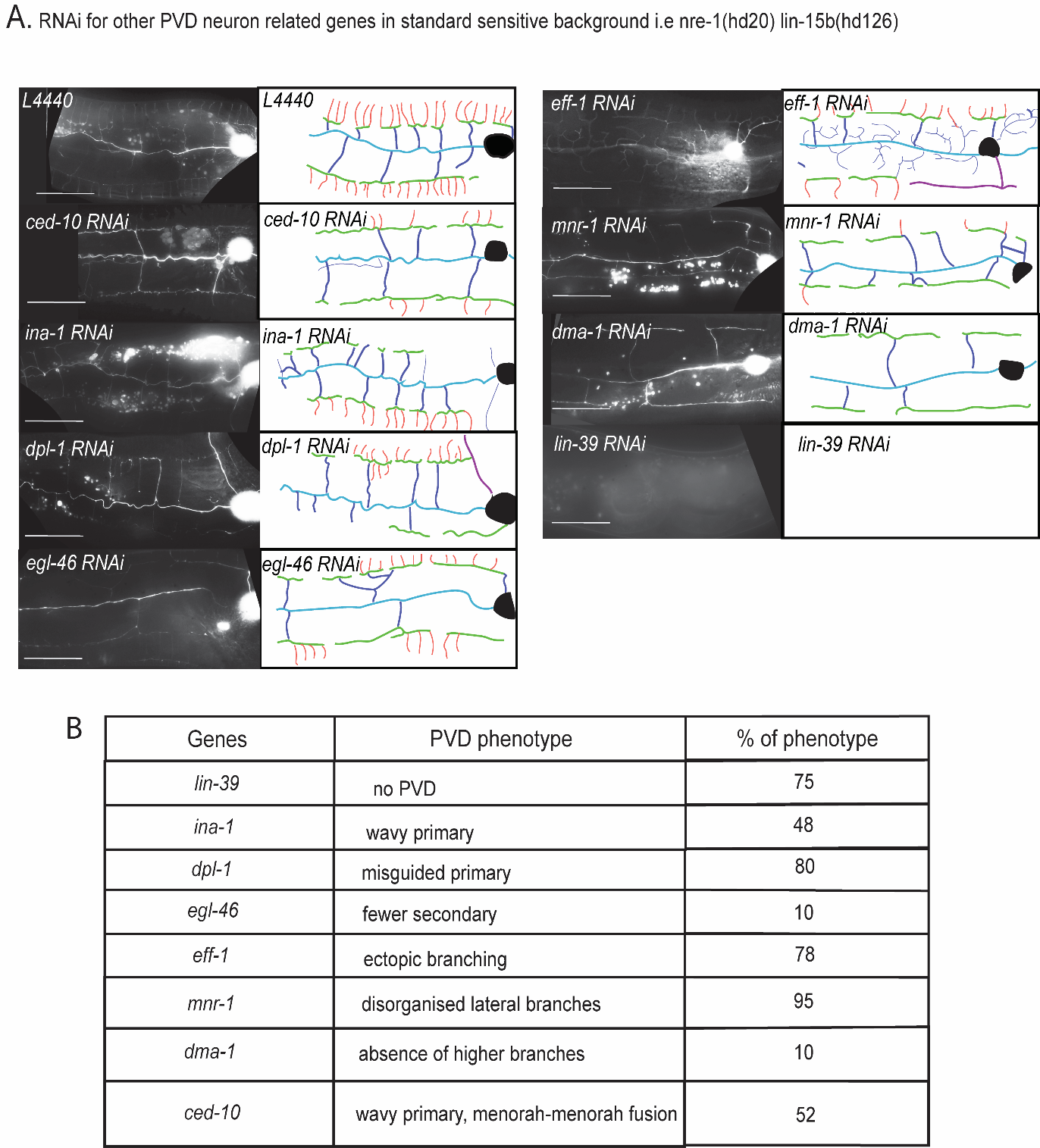

**Figure S2, related to Figure 1:**

(A) The images show the dendrite morphology defects in PVD neuron caused due to RNAi of various genes known to affect PVD development. The RNAi of these genes were performed in *nre-1(hd20)lin-15b(hd126)* background. The illustrations of the defects caused due to RNAi of these genes are also shown on the right. The hierarchy of PVD dendrite are shown in different colors i.e quaternary in red, tertiary in green , secondary in violet, primary in blue. Scale bar is 25 μm. (B) The RNA experiment mentioned in panel-A is summarized in a tabular form. The phenotypes associated to the RNAi of various genes are mentioned in this table. Biological replicates (1≤ N≤2).

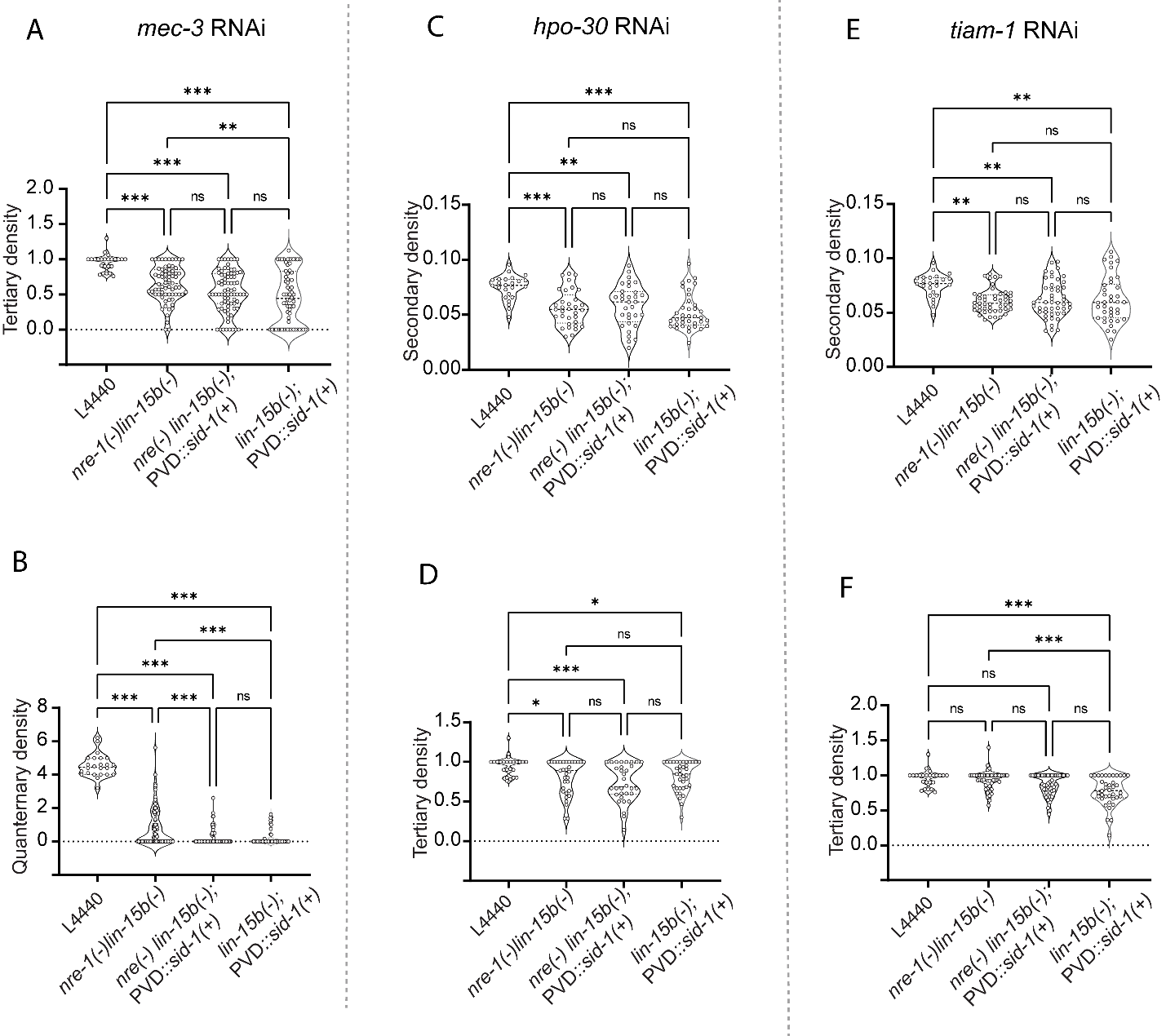

**Figure S3, related to Figure 3**

(A-B) Tertiary and quaternary density of PVD dendrites for *mec-3* RNAi in different sensitive backgrounds. (C-D) represents secondary and tertiary density of PVD dendrites for *hpo-30* RNAi worms. Similarly, (E-F) show the tertiary and secondary density for *tiam-1* RNAi. Biological replicates (1≤N≤4) and number of worms (24≤n≤97). Statistics: One-way ANOVA with Turkey’s multiple comparison test p<0.05*, 0.01**, 0.001***, ns (not significant).
